# A Kiosk Station for the Assessment of Multiple Cognitive Domains and Enrichment of Monkeys

**DOI:** 10.1101/2021.03.06.434198

**Authors:** Thilo Womelsdorf, Christopher Thomas, Adam Neumann, Marcus Watson, Kianoush Banaie Boroujeni, Seyed A. Hassani, Jeremy M. Parker, Kari L. Hoffman

**Author notes:** Corresponding Authors: Dr. Thilo Womelsdorf, Vanderbilt University, Psychology Department, 301 Wilson Hall, 111 21st Avenue South, 37240-1103 Nashville TN.

## Abstract

**Background:** Nonhuman primates (NHPs) are self-motivated to perform cognitive tasks on touchscreens in their animal housing setting. To leverage this ability, fully integrated hardware and software solutions are needed, that work within housing and husbandry routines while also spanning cognitive task constructs of the Research Domain Criteria (RDoC).

**New Method:** We describe a Kiosk Station (KS-1) that provides robust hardware and software solutions for running cognitive tasks in cage-housed NHPs. KS-1 consists of a frame for mounting flexibly on housing cages, a touchscreen animal interface with mounts for receptables, reward pumps and cameras, and a compact computer cabinet with an interface for controlling behavior. Behavioral control is achieved with a unity3D program that is virtual-reality capable, allowing semi-naturalistic visual tasks to assess multiple cognitive domains.

**Results:** KS-1 is fully integrated into the regular housing routines of monkeys. A single person can operate multiple KS-1s. Monkeys engage with KS-1 at high motivation and cognitive performance levels at high intra-individual consistency.

**Comparison with Existing Methods:** KS-1 is optimized for flexible mounting onto standard apartment cage systems. KS-1 has a robust animal interface with options for gaze/reach monitoring. It has an integrated user interface for controlling multiple cognitive task using a common naturalistic object space designed to enhance task engagement. All custom KS-1 components are open-sourced.

**Conclusions:** KS-1 is a versatile tool for cognitive profiling and enrichment of cage-housed monkeys. It reliably measures multiple cognitive domains which promises to advance our understanding of animal cognition, inter-individual differences and underlying neurobiology in refined, ethologically meaningful behavioral foraging contexts.

## 1. Introduction

Monkeys are housed in captive settings in zoos, primate service centers and research institutions. A rich, >30 years long history has shown that in these settings monkeys willingly engage in complex computerized cognitive tasks (Rumbaugh et al., 1989; Perdue et al., 2018). In their regular housing environments, monkeys (nonhuman primates, NHP’s) engage with joysticks or touchscreens, can semi-automatically train themselves on visual discrimination tasks, and when offered to freely choose amongst different tasks, they show motivation and insights into which cognitive tasks are most rewarding for them (Washburn et al., 1991; Gazes et al., 2013; Calapai et al., 2017; Fizet et al., 2017; Berger et al., 2018; Sacchetti et al., 2021). This prior work suggests a large potential to leverage the cognitive skills and the motivation of NHPs to (*1*) enrich animals’ cognition in their housing setting, (*2*) learn about their cognitive capacities and strategies to perform complex tasks, and (*3*) increase the ecological validity of brain-behavior coupling, through the concomitant use of species-typical, unrestrained behaviors (Lepora and Pezzulo, 2015; Krakauer et al., 2017; Datta et al., 2019).

The implementation of cognitively engaging tasks in captive settings faces several challenges. Chief among them is the difficulty to build the necessary hardware that fully integrates a touchscreen apparatus with the housing requirements. A second major challenge is the implementation of a cognitive task space for animals that meaningfully assesses performance across multiple cognitive domains. Here, we address both of these challenges.

Previous solutions of cage-based cognitive testing in animal housing environments provide guidance on how to build a cognitive testing apparatus adapted to animal cages (Washburn and Rumbaugh, 1992; Crofts et al., 1999; Weed et al., 1999; Mandell and Sackett, 2008; Fagot and Bonte, 2010; Nagahara et al., 2010; Truppa et al., 2010; Gazes et al., 2013; Berger et al., 2017; Calapai et al., 2017; Claidiere et al., 2017; Curry et al., 2017; Fizet et al., 2017; Tulip et al., 2017; Butler and Kennerley, 2018; Jacob et al., 2020; Griggs et al., 2021; Sacchetti et al., 2021). Whereas these tools have resulted in remarkable behavioral outcomes, their designs are not easily integrated in cage housing spaces, requiring e.g., a separate space for computer control or lacking an easy means to remove the tool during washing routines. They typically do not offer all desired features such as multiple camera mounts or options for fluid as well as pellet dispensers. Moreover, they vary widely in the validity and flexibility with which they assess different cognitive albitites. Many designs are not easily accessible on public repositories, have limited adaptability to incorporate improved experimental designs, and their advanced software packages are platform-dependent and may not entail a common cognitive task space that is desired for assessing multiple cognitive domains. Here, we propose an extension to existing approaches that addresses these challenges with a new, open-sourced variant of a touchscreen-based kiosk station (KS-1) for NHP’s.

The proposed KS-1 can be operated with any behavioral control suite; however, to address the second major challenge in adopting a touchscreen apparatus for cage-housed NHP’s we integrate KS-1 with an open-sourced control suite and document how a large common object space can be used in different tasks designed to assess multiple cognitive domains. Testing multiple cognitive domains is essential in clinical neuropsychiatric research because common disorders involve dysfunctions typically in more than one cognitive domain with common drug treatments affecting multiple domains (Knight and Baune, 2018; Zhu et al., 2019). For example, in major depressed subjects antidepressant drugs improve *executive function, attention and speed of processing, and learning/memory* domains (Harrison et al., 2016). In schizophrenia, too, multiple domains need to be considered., The MATRICS (*Measurement and Treatment Research to Improve Cognition in Schizophrenia*) consortium (Buchanan et al., 2005) proposes the MATRICS Consensus Cognitive Test Battery (MCCB) to measure multiple cognitive domains when assessing cognitive outcomes in treatment studies *<u>in schizophrenia</u>* (Buchanan et al., 2011). To address these criteria we document how the KS-1 can be used to routinely assess multiple MATRICS domains including *Speed of Processing, Attention, Working Memory*, and *Visual Learning* (Nuechterlein et al., 2004).

## 2. Materials and Methods

### 2.1. Subjects

Cognitive profiling and enrichment with cage-mounted kiosks was performed in six male and one female rhesus macaques (*Macaca mulatta*), ranging from 6-9 years of age and 8.5-14.4 kg weight. All animal and experimental procedures were in accordance with the National Institutes of Health Guide for the Care and Use of Laboratory Animals, the Society for Neuroscience Guidelines and Policies, and approved by the Vanderbilt University Institutional Animal Care and Use Committee.

### 2.2. Hardware and setup

The kiosk consists of two modules that are easily connected and disconnected from each other: (1) a “front-end” arcade interface for the animal that connects to a mounting frame on the cage, replacing one of the cage’s side panels, and (2) a “back-end” cabinet for hardware and hosting a user interface (**Fig. 1**, technical details in **appendix 5.1**, resources available at https://github.com/att-circ-contrl/KioskStation). The kiosk replaces the front panel of an apartment cage and provides a 19.5’’ touchscreen within reach from the front of the cage. The front-end (facing the animal in the cage) is a robust stainless-steel enclosure with a receptacle for pellet rewards, a sipper tube for fluid reward, three plexiglass shielded window openings for cameras, a window opening with a lockable door to allow personnel reaching from outside in (for cleaning), a cut-out for the touch screen (mounted in the back-end but flush with the front-end when assembled), and a plexiglass window below the screen for eye and head tracking devices. A reward pump and pellet dispenser are mounted outside at the side of the front kiosk part. The back-end cabinet of the kiosk is secured to the front-end using mounting pins, two slide bolts, and two machine screws, for ease of assembly and disassembly. A similar arrangement secures the front end to the mounting ring. The back-end hosts the touchscreen, the experiment computer, the camera control computer, a wireless router, various auxiliary equipment described below, a small monitor, keyboard, and trackball mouse that the operator/trainer/experimenter uses for experimental control and animal monitoring (**Fig. 2**). The touchscreen is enclosed in a rigid aluminum shell designed to provide a robust interface for sustained animal interactions. The back-end cabinet’s shelves can be arranged flexibly and loaded with custom equipment, with cable ports providing access to equipment mounted on the front-end and two fan ports with air filters providing cooling for electronics. An overview of kiosk construction and contents is shown in **Fig. 1** and **Fig. 2**, and a list of kiosk-related equipment is provided in the **appendix 5.1**.

**Figure 1.**
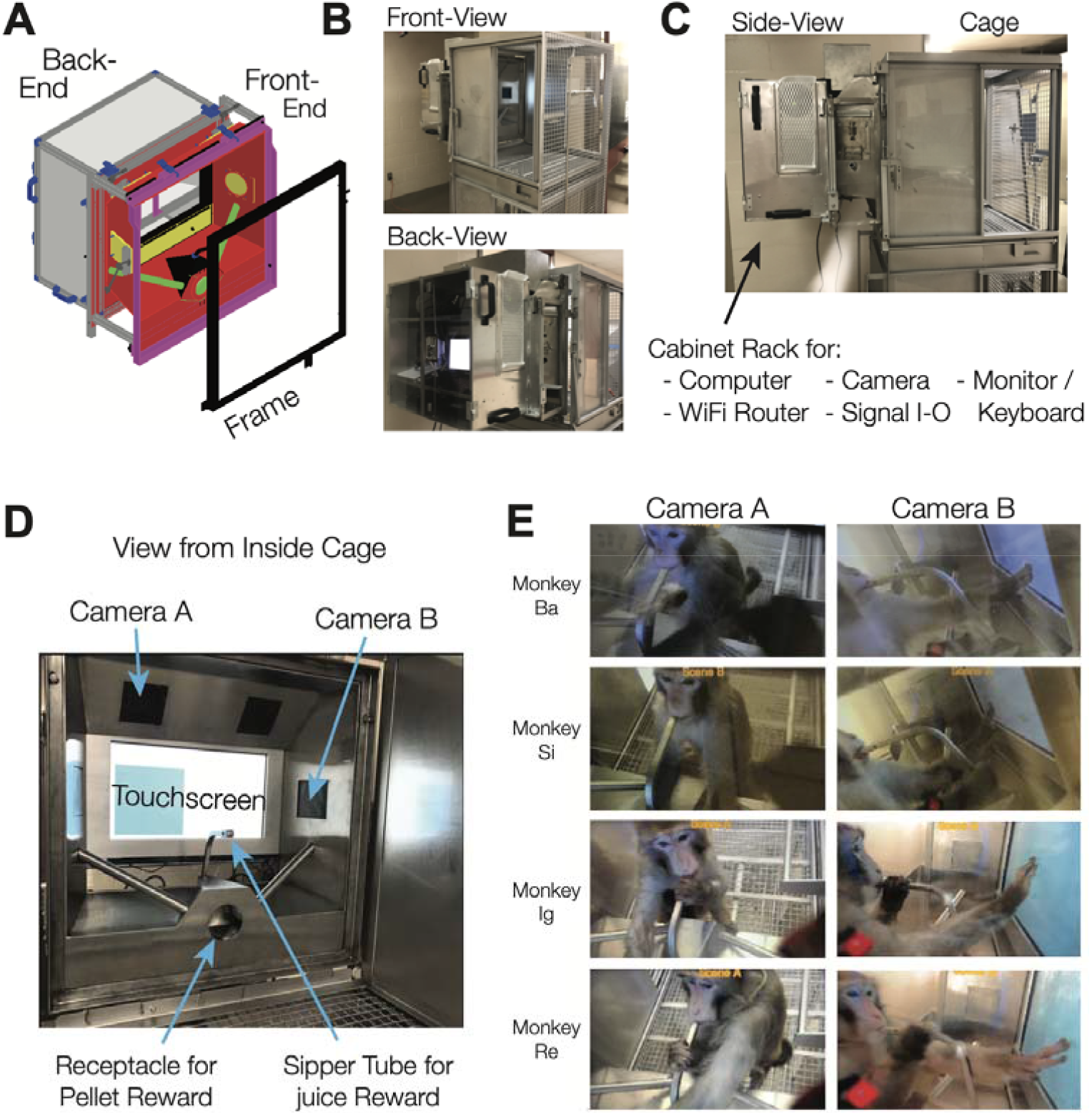
Kiosk design and cage-mount. (**A**) Design drawing of the frame, front-end, and back-end of the Kiosk. The design is available as cad file online (see appendix). (**B**) Front- and Back-end of the Kiosk Station mounted to a Primate Products Inc. apartment cage for rhesus monkeys. Side-view of the mounted Kiosk which extends ∼23” from the cage. The side views shows a small locked side door at the front end that enables reaching inside for cleaning the touchscreen. inside view onto the interface to the monkey shows the touchscreen, camera windows, sipper tube and pellet receptacle. (**E**) Four monkeys (rows) interacting with the touchscreen as seen from the top and side camera window (camera windows A and B in panel *D*). All monkeys maintain mouth contact with the sipper tube awaiting fluid reinforcement for their behavior (*left*) and use their fingers to touch objects displayed on the screen (*right*).

**Figure 2.**
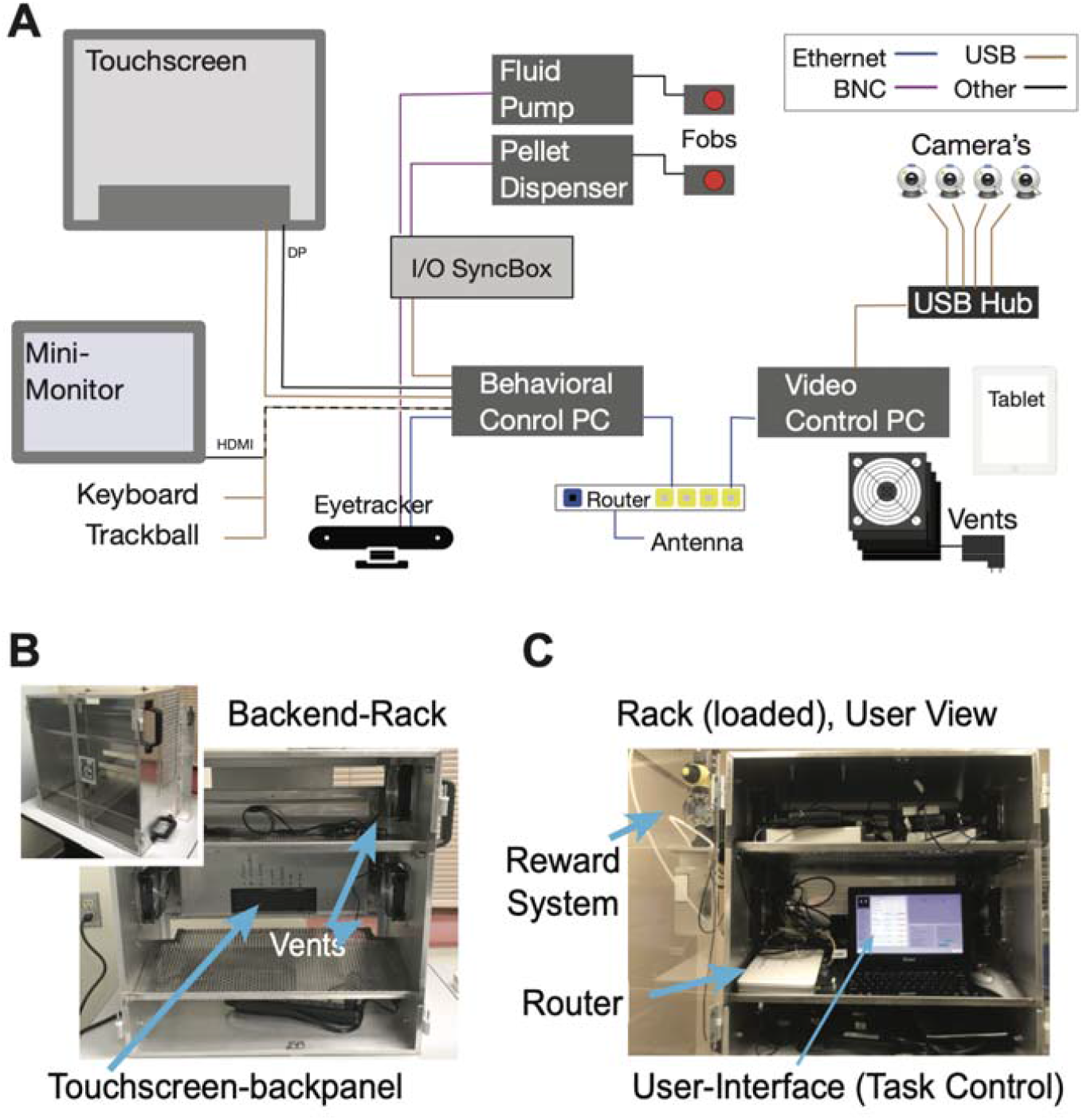
Electronic hardware organization. (**A**) Diagram showing cable connections between the main electronic Kiosk components. (**B**) The back-end of the Kiosk is a rack with three levels. It has a transparent door, an aluminum frame for the touchscreen, usb-powered vents on both sides, and handle-bars mounted outside of the rack. (**C**) View inside a hardware loaded Kiosk rack. A 11” monitor, keyboard and trackball mouse sit on an angled shelf allowing personnel to control behavioral tasks, video-streaming and reward delivery.

### 2.3. Video monitoring of animal behavior

The kiosk front-end contains plexiglass windows for camera surveillance of the monkey’s performance. There is a window for a side view, two windows for top-down views on to the monkey and a large horizontal plexiglass window below the touchscreen for a bottom-up view of the monkey’s face and shoulders for tracking gaze and arm movements. Each window contains mounting adapters for cameras. We typically use Logitech C930e digital cameras which have 90°-100° field of view (FOV). A custom-built multi-camera streaming system (*NeuroCam*) controls and synchronizes up to five cameras (see **appendix 5.1**, technical details and firmware available at <https://github.com/att-circ-contrl/NeuroCam>). The *NeuroCam* has a web browser interface to configure each camera’s resolution and frame rate and to monitor up to five cameras simultaneously. The NeuroCam control computer is located in the kiosk’s back-end cabinet and can be securely accessed by external wireless devices (e.g. tablets or smartphones) (**Fig. 2C**). This allows monitoring the animals task engagement in the kiosk from outside the housing room and the recording of up to five synchronized camera views which will allow 3D reconstruction of gaze and reach patterns (Karashchuk, 2019; Sheshadri et al., 2020).

### 2.4. Power allocation

The power requirements of the kiosk station can be met with a single regular power outlet. For the KS-1 in Fig. 1 and Fig 2, the behavioral control computer was a NUC8i7HVK with a 230W power supply and the video control computer was a NUC6i7KYK with a 120W power supply. The remaining equipment uses approximately 80W (ELO 2094L touchscreen: 20W; Eyoyo 12” monitor: 24W; Asus RT-N66U gateway router: 30W; four Sunon EEC0251B3-00U-A99 fans: 9W). Cameras are USB-powered through the camera control computer, and the reward pump and pellet dispenser have only low and transient power consumption, so they do not contribute significantly to the total power requirements. For the configuration described, the KS-1 hardware consumes approximately 430W.

### 2.5 Software suite and behavioral control

The kiosk can be run flexibly by any behavioral control software that registers touchscreen interactions and controls the reward delivery to the animal. Here, we propose using the *Unified Suite for Experiments* (*USE*) (Watson et al., 2019b), which is an open-sourced suite of C# scripts that extend the Unity video game creation engine (Unity-Technologies, 2019) to be a robust experimental design and control platform. USE can run multiple visual-cognitive tasks using different response modalities (touch, joystick, buttons, gaze) and reward feedback delivery types (primary fluid/food and secondary vis./audit. rewards) while being fully integrated with a I/O system that allows communication, control and synchronization of time stamps with experimental hardware (e.g. eyetrackers, reward systems, wireless recording devices). Unity 3D and USE are platform independent with any modern computer. The programming code of USE is freely available and a documentation and user manual are available online (see **appendix 5.2** and https://github.com/att-circ-contrl/use) (Watson et al., 2019b). Although USE can be customized by users with programming expertise, no computer programming is needed to run various conventional cognitive tasks.

USE enables experimental control at multiple granularities, from individual trials or blocks, to the task as a whole. Text files controlling parameters at each of these levels can be generated as needed. Thus, for some tasks, *trial definition* configuration text files are used to control the specific stimuli shown on each trial, their precise size, positions and orientations, and the reward magnitude and probabilities associated with each. For others, we use *block definition* files, that define rules governing reward on each task across many trials, and the suite uses these rules to choose and display appropriate stimuli, without the need for the user to specify the details for individual trials. For others, we use a mix of the two to enable lower- and- higher-level control over different aspects of the experiment as needed.

During an experimental session, USE controls both the display shown to the participant and a separate display shown to the operating personnel. At the start of a session, the operator’s display enables the selection of the desired configuration files, the path at which data will be saved, paths at which stimuli are stored, and so on. During the remainder of the session, the operator’s display includes information summarizing participants’ current performance, a set of sliders and buttons that enable real-time control over aspects of the experiment (e.g. inter-trial interval duration, or distance thresholds for gaze or touch to be considered as on an object), and a window that mirrors the participant display, with overlaid information such as gaze traces, touch locations, or highlights over particular stimuli (**Fig. 2C**).

USE saves data for each individual frame, enabling complete reconstruction of the entire experimental session, if needed. Data is saved after each trial to allow termination of ongoing task performance without loss of data. A set of MATLAB scripts are available as an online resource to preprocess data into an efficient format for analysis and visualization (see **appendix 5.2**).

### 2.6 Unified multidimensional object set

During training, the animals are adapted to a large set of 3D-rendered objects having multiple feature dimensions (Watson et al., 2019a). This ensures animals are pre-exposed to all the visual features of objects that will be used as target or distractor features for cognitive tasks after initial training is completed. The large feature space provided by multidimensional *Quaddle* objects described by Watson and colleagues (Watson et al., 2019a) is pre-generated and integrated in the USE behavioral control suite. Each Quaddle object has a unique combination chosen from nine body shapes, eight colors, eleven arm types and nine surface patterns, providing 7128 (9×8×11×9) unique objects. The objects are generated with customizable batch scripts for the software Autodesk Studio X Max and are available online (http://accl.psy.vanderbilt.edu/resources/analysis-tools/quaddles/). For the cognitive task object colors are selected to be equidistant within the perceptually-defined CIELAB color space. Typically, the objects are rendered to extend ∼1-2’’ on the screen and are presented on an Elo 2094L 19.5 LCD touchscreen running at 60 Hz refresh rate with 1920 x 1080 pixel resolution.

### 2.7. Kiosk training procedure

Before animals perform complex cognitive tasks, they undergo a training regime that standardizes their touch behavior and ensures pre-exposure with all visual object features used in later cognitive task variants. For all training steps, the animals are given free access to the kiosk for 90-150 minutes per day irrespective of the time they engage with the touchscreen. In the first training step animals learn to touch an object extending ∼1-2” on the screen at random locations, hold the touch for 200-300 ms, and release the touch within 500ms in order to receive a reward feedback. This Touch-Hold-Release (THR) task proceeds through pre-defined difficulty levels that the operator/tester can set flexibly before or during task performance. Initially, the animals receive reward for touching a large blinking blue square, which successively gets smaller and is presented at random locations to train the precision of touching a blue square in its immediate perimeter. In parallel with training touch precision, reward is provided upon touch release (as opposed to onset), and the minimum and maximum durations for touching the object to receive reward is standardized. Animals move through these difficulty levels until they are considered “touch-ready”, similar to the ‘joystick-ready’ criterion successfully used in the context of the ‘Rumbaughx’ (Perdue et al., 2018), which in our experience occurs within ∼2-6 weeks. We had three of seven animals temporarily showing suboptimal, undesired touch behavior such as swiping or briefly tapping the screen instead of showing precise touches of an appropriate duration. THR training gradually eliminates such suboptimal strategies.

In the second training step animals learn the detection and discrimination of more complex objects by choosing one among several visual objects on the screen, with one being rewarded. This visual search task proceeds through increasing difficulty levels. Trials are started by touching a central blue square. Then a target Quaddle is shown in the presence of 0,3,6,9 or 12 distracting Quaddles. The easiest difficulty level is a feature popout visual search task in which a target object is distinguished from distractor objects by one visual feature. Quaddle objects are rendered with features from a common multidimensional feature space consisting of different arms types, body shapes, surface pattern and color (see above). A single set of features within this feature space will never be rewarded. These never rewarded, or ‘neutral’ features include a grey color, uniform surface pattern, spherical body shape, and straight blunt arms. Touching a Quaddle with all four neutral features aborts a trial without reward, thus incurring a temporal delay, or cost, for the animal before initiating the next trial. At later difficulty levels, the target Quaddle has non-neutral features in more than one visual dimension, e.g. having a unique color, surface pattern and arm type, but still the ‘neutral’ spherical body shape. This target object is then presented together with distracting objects that also have non-neutral features in one, two or three dimensions (**Fig. 3A**,**B**). The number of feature dimensions varying in distractor objects determine the amount of interference animals experience during visual search for the target. Upon completion of the second training step the animals are therefore able to perform top-down visual search in the presence of up to twelve distractor objects and targets sharing features with distractors in up to four feature dimensions.

**Figure 3.**
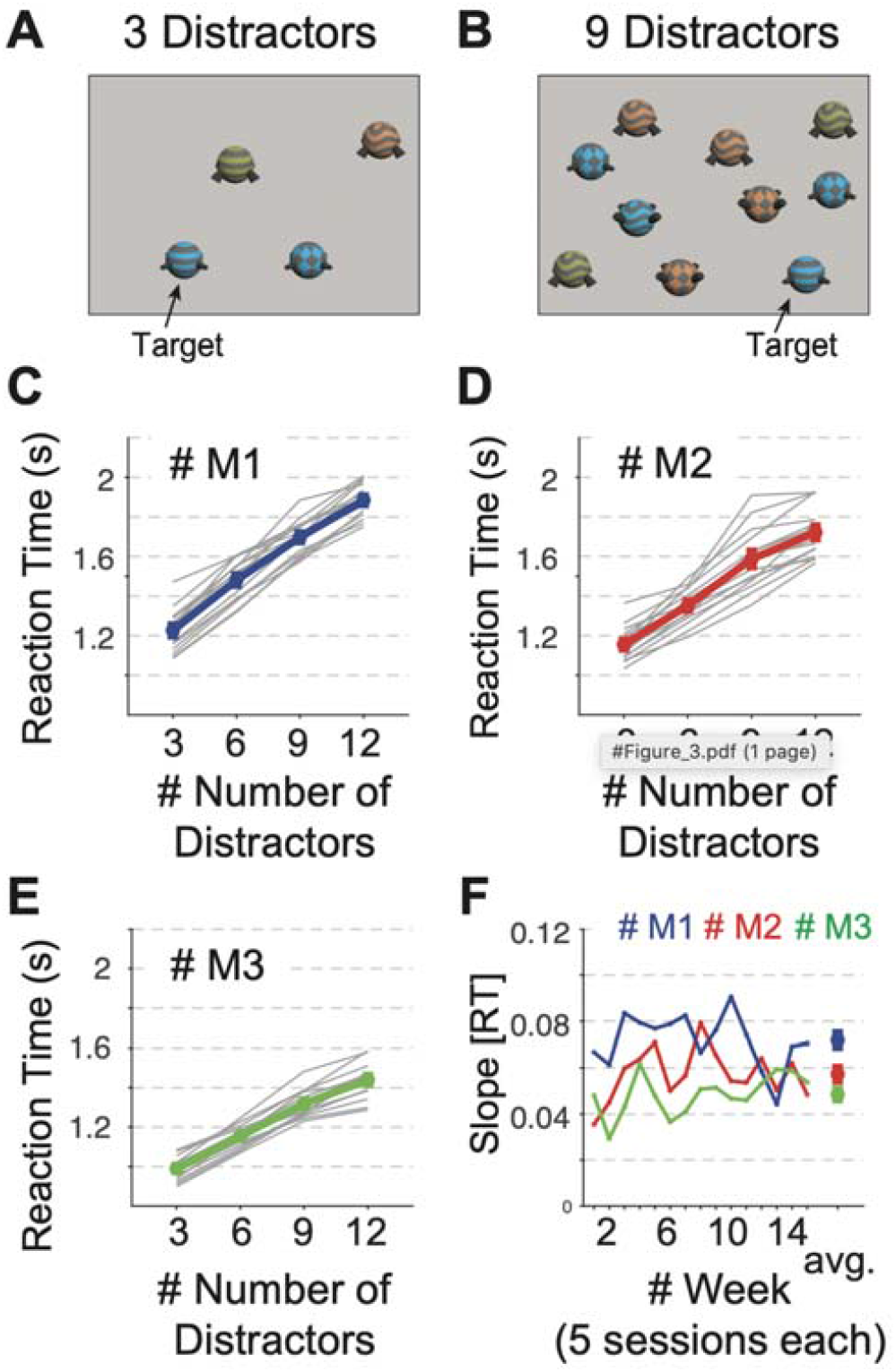
Visual search performance. (**A, B**) Visual display with a target object and 3 (*A*) and 12 (*B*) distractor objects that shared features with the target. (**C, D, E**) Visual search reaction times (Y-axis) for 3-12 distractors (x-axis) in each of 15 weeks (grey lines) and on average (colored line) for monkey M1 (C), M2 (D) and M3 (E). (**F**) The regression slope or the reaction time increase with distractors for each monkey (in color) over 15 weeks. The rightmost data point is the average set size effect for each monkey. Error bars are SEM.

The third training step extends the task competencies of the animals to the domain of cognitive flexibility. Cognitive flexibility is measured with a feature-value learning task by how quickly and accurately animals adjust to changing reward contingencies in their environment. We test cognitive flexibility using displays with 3 or 4 objects among which only one object contains a rewarded feature. The kiosk training regime indexes cognitive flexibility by changing the rewarded object feature every 40-60 trials and measuring how fast the animals adjust their choice to the newly rewarded and away from the previously rewarded feature. This flexible feature rule learning task is trained at different difficulty levels. At the easiest difficulty level, the animals are presented on each trial with three objects that have different features in only one feature dimension (e.g. their arms might differ), while all other dimensions have neutral features (**Fig. 4A**). Only one feature value is rewarded, thus creating a 1-dimensional, or 1-way learning problem. At later stages of learning target and distractor objects vary features in two or three dimensions (**Fig. 4B**). This variation creates a 2-way and 3-way feature space that the animals need to search to find the rewarded target feature, i.e. it creates a learning environment with parametrically increasing attentional load.

**Figure 4.**
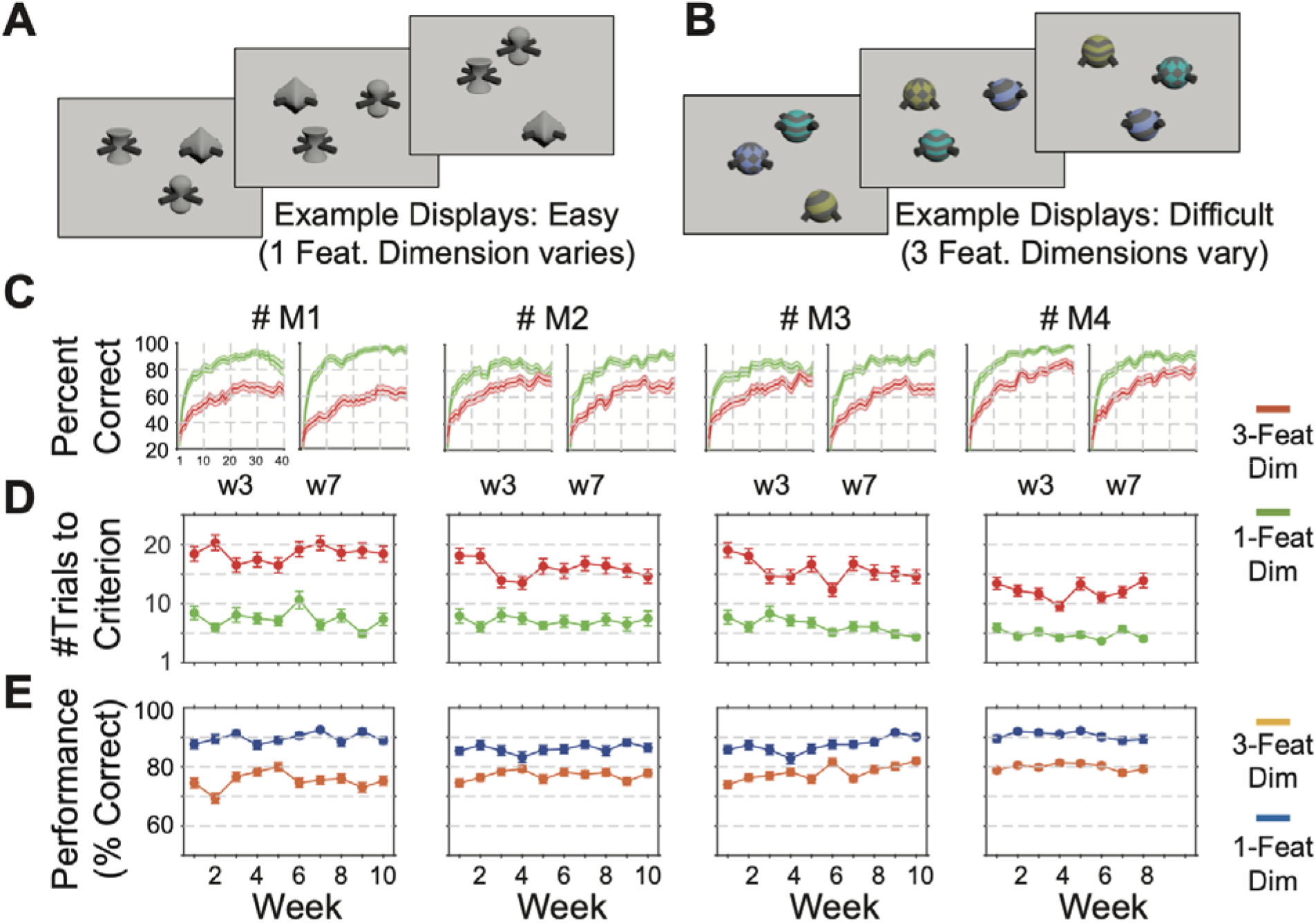
Measuring cognitive flexibility. (**A, B**) Example displays with three multidimensional objects for the feature-reward learning task. The task required monkey to learn which feature is linked to reward in blocks of 45-55 trials. The target object that was rewarded was defined by a specific feature. Objects varied trial by trial either in features of one dimension (e.g. only body shapes in *A*) or of two or three dimensions (e.g. pattern, color and arm types in *B*). (**C**) Performance learning curves averaged over five sessions in weeks 3 and 7 (w3, w7) for monkeys 1 to 4 (*left* to *right*). Green and red lines correspond to the easy and difficult condition (varying 1 or 2/3 feature dimensions. (**D**) For four monkeys (columns) the average number of trials needed to reach learning criterion in the easy and difficult conditions over 15 weeks (5 sessions per week). (**E**) Same format as D for the plateau performance of monkeys in trials after the learning criterion was reached. Error bars or shading correspond to SEM.

### 2.8. Task structure for visual search and feature rule learning

We tested the performance of monkeys in the kiosk environment with a visual search task that varied the target-distractor similarity (1 to 3 target features shared among distractors), and with a flexible feature learning task that varied the number of interfering object features (1 to 3 feature dimensions varied). For both tasks a trial started by presenting a blue square in the center of the screen. When touched for 0.2 s the square disappeared and after a 0.3-0.5 s delay the task objects were presented. In the visual search task there were always 10 initial trials in which the same object was presented alone for up to 5 s or until the monkeys touched it for at least 0.2 s, which triggered visual and auditory feedback and the delivery of fluid reward though the sipper tube. These ten *initialization* trials defined the target object for subsequent trials in which the target was shown together with 3, 6, 9, or 12 distracting objects randomly at intersections of a virtual grid. The number of distractors varied randomly over a total of 100 search trials. In each experimental session the animals performed the visual search task twice (each with 10 initialization trials and 100 test trials). Other tasks not discussed here were run during the same sessions.

In the feature learning task, 0.5 s after the offset of the central blue square, three multidimensional objects appeared at random intersections of the grid locations. Objects spanned ∼1.1” in diameter and were 4.1” away from the display center. One of the three objects contained a feature that was associated with reward (e.g. the oblong shape) for a block of 45-60 trials, while there was no reward associated with other feature of the same feature dimension (e.g. spherical or cubic shapes) or of other feature dimensions (different colors, arm types or surface patterns). The monkeys had up to 5 s to choose one of the objects by touching it for >0.2 s, which triggered visual and auditory feedback. If the chosen objects had the rewarded feature it was also followed by the delivery of fluid reward. Each experimental testing session presented 40 learning blocks, in which the target feature dimension and feature of that dimension was defined randomly among four possible feature dimensions (shape, color, arm type, or body shape) and among 7-11 different feature values (e.g. different colors, different arm types) of that dimension.

### 2.9 Testing procedure

All testing proceeded in cage-mounted kiosks in the housing rooms. The kiosk was mounted to one apartment cage unit. For the duration of testing that apartment cage and the neighboring apartment cage were both freely accessible to a single rhesus monkey. Before and after the behavioral sessions, the monkeys were pair-housed and only separated for the duration of the kiosk performance sessions which lasted 90-120 min with rare exceptions with longer duration. For each monkey, the fluid reward (water) volume was adjusted such that the completion of the task would provide between ∼150-350 ml of water, corresponding to ∼20-25ml/kg for individual animals. All monkeys would work for more fluid reward on some days, but this would then result in reduced motivation on the following days, evident in a reduced number of completed learning blocks and reduced performance levels. Without fluid control the monkeys engaged with the kiosk tasks, but made pauses during task engagement, which was quantifiable in overall lower performance and inconsistent performance.

### 2.10. Behavioral analysis

Behavioral data generated within the KS-1 are preprocessed with scripts written in MATLAB (The Mathworks, Inc.). They are openly available with the USE experimental suite (https://bitbucket.org/MarcusRWatson/use_analysis). Visual search performance was analyzed per session and then pooled across five sessions of a week. The average reaction times at increasing number of distractors (3/6/9/12) were fit with a linear regression to estimate the slope (indexing the set size effect) and the intercept (indexing the baseline reaction time speed), similar to previous studies (Purcell et al., 2010; Purcell et al., 2012). Cognitive flexibility during the feature-reward learning task was evaluated as the number of trials animals needed within each block to reach criterion performance of 70% correct choices over ten trials as in previous studies (Hassani et al., 2017). Plateau performance was calculated as the accuracy across trials following the trial at which the learning criterion was reached to the end of the block.

## 3. Results

### 3.1. Kiosk placement and handling

The KS-1 is mounted to regular housing cages of pair-housed rhesus monkeys, replacing the front panel of the cage (**Fig. 1A, B**). Operating the kiosk is integrated with the regular husbandry protocols and with the enrichment programs for the animals. The kiosk has a small footprint, extending 23” inches from the cage into the room (**Fig. 1C**). The distance of the kiosk cabinet to the wall or cage opposite of the kiosk mounted cage is 38-44” inches. With these dimensions it does not interfere with cleaning routines inside the housing room and is easy to operate by standing in front of it. When the cage needs to be moved to e.g. a cage-washer, a single person can unmount the fully-loaded kiosk from the cage and safely place it on a table-lift trolley for temporary storage (**appendix 5.1**). The kiosk unmounts by unlocking two hooks from the cage frame and loosening a separation screw at the bottom of the kiosk. No cables or electronics need to be changed when unmounting or mounting the kiosk. The ease of handling is achieved through the modular design with the kiosk’s front-end being locked into the kiosk’s cage frame with spring loaded hooks. The KS-1’s described here are removed bi-weekly during cage washing.

### 3.2. Effectiveness of animal interface

The KS-1 front end provides the interface for the animal with a 19.5’’ touchscreen embedded in an aluminum frame and recessed ∼11’’ away from the cage border (**Fig. 1D**). A stainless**-**steel sipper tube protrudes from the center console up towards the animal at a height and distance from the screen that can be flexibly adjusted to the optimal position for monkeys of different sizes with screws below the kiosk. When engaging with the touchscreen, animals generally make contact with the sipper tube’s mouth-piece so that it serves as a means to control the distance of the animals to the touchscreen (see the typical positioning of four monkeys in **Fig. 1E**). Rhesus monkeys of approximately ≥3 years of age will be able to reach to and touch all corners of the touchscreen without moving their body away from the central spot in front of the sipper tube. The center console also has a receptacle into which pellets can be released from outside through ‘sliding’ tubes protruding from the outside into the center receptacle from the sides of the kiosk.

The front-end of the kiosk also provides three windows for cameras in the top-panel and in one-side panel. The other side panel has an opening window-gate allowing the operator to easily reach into the kiosk and clean the touchscreen when needed (visible in **Fig. 1C**). Below the touchscreen is a plexiglass panel that allows a free field of view from outside towards the space that contains the head (for face and gaze analysis) and shoulders of the animals. This is useful for eye- and body-tracking systems (Mathis et al., 2018; Karashchuk, 2019; Bala et al., 2020; Sheshadri et al., 2020).

### 3.3 Effectiveness of user interface

Operating the kiosk is accomplished through a computer and monitor interface located on the cabinet of the back-end of the kiosk. The kiosk back-end is a ventilated cabinet with electronic hardware including a computer running the touchscreen-based tasks, a computer streaming multiple cameras, a router with external antennas for fast WIFI access, an input-/output-box controlling peripheral devices (e.g. reward pump), as well as a user interface with a 12’’ monitor, keyboard and a trackball mouse (**Fig. 2A**). Hardware details are listed in the **appendix 5.1**. The hardware and cabling inside the cabinet can be spatially arranged at three shelving levels (**Fig. 2B**). A loaded cabinet is shown in **Fig. 2C**. It has the monitor and keyboard inside the cabinet at a height of ∼4.2’ from the ground allowing easy access to personnel standing in front of it. The transparent opening doors facilitate quick checking of the modus of operation while walking by the kiosk. This back-end user interface allows controlling all aspects of the task performance including the video monitoring of the monkey inside of the kiosk. The current installation has a remote control that allows manually controlling the opening of the reward pump or pellet dispenser to probe the animals motivation to approach the sipper tube for reward or test the reward systems functionality. The reward pump and pellet dispenser are mounted rigidly to the outside frame of the kiosk’s front-end (see **Fig. 2C**).

### 3.4 Software control

There are many systems that could control behavior in the kiosk, register behavioral responses and elicit TTL pulses for opening fluid/food dispensers to reinforce the behavior (Brainard, 1997; Peirce, 2008; Eastman and Huk, 2012; Doucet et al., 2016; Hwang et al., 2019). We operate the kiosk system with a custom developed, freely available, open-source software called *Unified Suite for Experiments* (USE) (Watson et al., 2019b). USE is integrated with an input-output box (‘I/O Synchbox’) for communication with reward systems and temporal synchronization with other devices such as video cameras and eye trackers. The Arduino based hard- and firmware of the I/O Synchbox are available and the set-up is documented (**appendix 5.1**, https://github.com/att-circ-contrl/SynchBox). USE is built on the unity3D video gaming platform to allow the use of 3D rendered objects and scenes in behavioral tasks, which are experimental options that have been shown to enhance the degree of engagement with touchscreen behavior (**Fig. 2A**) (Bennett et al., 2016).

USE provides additional features facilitating kiosk cognitive training and testing. Upon startup, USE shows a graphical user interface for selecting specific files and folders that contain the task protocol, the timing and calibration parameters needed, the path to visual objects and to data folders. This pre-selection eases use of the same kiosk with different animals that perform different tasks or require different task configurations. During task performance users can monitor the monkey’s performance online through a thumbnail image duplicating the front-end touchscreen display. The display overlays information about which object is rewarded and shows touchscreen touches of the monkeys with a history trace on the duplicated image display to allow tracking monkey’s choices (**Fig. 2C**). The user interface also allows the user to adjust multiple task parameters online during task performance such as the timing of the stimuli or inter-trial intervals, the required hold duration for registering touches, or the difficulty level for semi-automated early stages of training animals to touch, hold on, and release touch after a hold-duration of 0.1-0.4 s.

### 3.5. Cognitive profiling

The kiosk station and its integrated USE program allows profiling higher cognitive functions. We documented the consistency of profiling with a visual search task to quantify attentional filtering abilities, and a feature-rule learning task to quantify cognitive flexibility of the animals.

#### Visual search performance

We found that all three monkeys trained and tested in the kiosk on a visual search task showed the classical set-size effect of slower choice reaction times with increasing numbers of distractors (**Fig. 3**). The task cued a complex target object by showing it alone on the screen in ten trials. Thereafter, the target was embedded in displays with 3, 6, 9 or 12 distractor objects presented at random intersections of a grid spanning the touchscreen. Distractors shared one to three different features with the target making this a conjunctive-feature search task at different difficulties. Across 82, 72, and 74 testing sessions for monkeys M1, M2, and M3 respective we observed high average accuracies of >80% for all monkeys. M1, M2, and M3 detected the target at 87.5% (STD: ±6.98), 90.2% (±9.80), and 81.2% (±9.83), respectively. We grouped the first 15 weeks of kiosk sessions (5 sessions per week) and found reliable set size effects for each monkey at all times (**Fig. 3 B-D**). The regression slopes indicated that monkeys differed in their visual search performance. The highest distractibility was found for monkey M1 (slope 0.072 ±0.012, range [0.036-0.079]), followed by M2 (slope: 0.057 ±0.011, range [0.036-0.079]) and M3 (slope 0.049 ±0.009, range [0.029-0.062]). These slopes reflect that target detection was slowed on average by 72, 57, and 49 msec for each added distractor for monkeys M1, M2 and M3, respectively. The low standard errors of the slope estimates illustrate a high intra-subject reliability of attentional filtering abilities of the animals. The same rank ordering of monkeys was evident in their average baseline detection response time, or speed of processing, indexed as the intercept of the regression fit to the set size, with 905 (±99), 879 (±96), and 770 (±56) msec for monkeys M1, M2, and M3.

#### Cognitive flexibility

To test whether the kiosk environment allows reliable estimation of cognitive flexibility we trained and tested four monkeys on a flexible feature-reward learning task that varied attentional demands. In blocks of 45-55 trials the animals had to learn through trial-and error which object feature is consistently rewarded. The target object and two distractor objects varied either in 1, 2 or 3 feature dimensions, which increased the task difficulty by increasing the uncertainty about which feature was linked to reward and which features were unrewarded (**Fig. 4A, B**). We tested learning flexibility in 228 experimental sessions (63 sessions in monkeys M1, M2 and M3 and 39 sessions in M4). Sessions lasted 70-120 min during which monkeys completed all 40 learning blocks that were provided in the largest majority of sessions (avg. number of completed blocks per sessions: 39.9). Each monkey showed reliable learning curves across the whole testing period. Example learning curves for sessions in weeks 3 and 7 are shown in **Fig. 4C**. Monkeys showed consistent performance over 10 weeks but differed from each other. Learning speed, indexed as the average trial needed to reach 75% criterion performance over 10 trials, yielding on average, for monkeys M1-M4: trial 12.1 ±1.4 (range [9.6-13.9]), 15.9 ±1.6 (range [13.5-18.2]), 15.7 ±2.0 (range [12.3-19.1]), and 18.5 ±1.3 (range [16.5-20.4]). Plateau performance for trials after learning criterion was reached in individual blocks for monkeys M1-M4 of 75.3% ±2.9 (range [69.3-80.0]), 77.1% ±1.6 (range [74.5-79.2]), 78% ±2.7 (range [73.9-82.0]), and 79.9% ±1.2 (range [77.9=81.4]), respectively. The low standard errors for the monkey specific learning speed and plateau performance indicates a high intra-subject consistency.

### 3.6. Observations about animal kiosk engagement

Animals engaged with the Kiosk whenever it was made available to them, and showed consistent motivation to engage with the Kiosk for prolonged periods of time. Typically, an animal waits already at the gate of the apartment cage before it is opened by the operator with the gate remaining open for 90-120 minutes on weekdays daily so that animals can choose whether to engage with the kiosk. This indicates anticipation and motivation to engage with the Kiosk touchscreen and confirms prior reports that Kiosk engagement is a form of cognitive enrichment (Washburn and Rumbaugh, 1992; Bennett et al., 2016; Calapai et al., 2017; Egelkamp and Ross, 2019). There are few exceptions to this behavior. One animal took more time to engage with task initiation at a time when the amount of dry biscuits was reduced for dietary reasons suggesting that animals are more motivated to work for fluid reward when they have a regular dry food diet available at the time or prior to engaging with the task. Moreover, two animals took breaks half-way during their 90-120 min sessions and walked into the second apartment cage to pick up chow or produce before continuing task engagement.

## 4. Discussion

We have documented an open-sourced hardware and software solution for the cage-based assessment of multiple cognitive domains and the cognitive enrichment of rhesus monkeys. We validated multiple KS-1, each providing 2 pair-housed animals daily sessions of cognitive enrichment and assessment. The animal interface enables animals to engage with cognitive tasks for rewards in a controlled and stereotyped way providing reliable, high quality cognitive-behavioral performance data. Its hardware is fully integrated with a software suite for temporally precise behavioral control and a video monitoring device for high resolution animal tracking. Hard- and software components can be handled professionally by a single person with little training. All custom designed hard—and software components are open-sourced supporting easier adaptability of the integrated software (White et al., 2019) (**appendix 5.1**).

### Enrichment and assessment of multiple cognitive domains

We have shown that the KS-1 succeeds to cognitively engage monkeys over multiple weeks. Such a computer based cognitive engagement is considered a versatile cognitive enrichment strategy that can effectively promote the psychological well-being of NHPs (National-Research-Council, 1998) (see also: The Macaque Website: https://www.nc3rs.org.uk/macaques/ hosted by the National Centre for the Replacement, Refinement and Reduction of Animals in Research (NC2R) in the UK). The cognitive assessment of monkeys over 10 weeks with a flexible learning task and over 15 weeks with a visual search task resulted in means and low standard errors of performance scores that distinguished different monkeys and showed high consistency within individual monkeys. Such intra-individual stability is typically interpreted as indexing strong cognitive ability of individual subjects, and offers the sensitivity to distinguish among subjects (Slifkin and Newell, 1998). These results suggest the KS-1 can serve as a tool to assess inter-individual cognitive differences between NHP’s and to track their changes over the lifespan and across different experimental conditions. The behavioral data we presented further document that this assessment can include multiple cognitive domains. These domains include multiple constructs of the RDoC Matrix that serves as diagnostic guide for the understanding of dysfunctional brain systems underlying psychiatric diseases (Cuthbert and Insel, 2013). The visual search task we used measures not only set size effects that indexes the efficiency of attentional filtering of distraction. It also quantifies the speed of processing (baseline search speed) that is a known behavioral marker of aging. Visual search tasks are easily extended to obtain indices for multiple other domains including, for example, indices of perceptual interference by varying the target-distractor similarity, or to obtain indices of reward-based capturing of attention by varying the expected value of distractors (Wolfe and Horowitz, 2017; Wolfe, 2021).

Similarly rich in opportunities to quantify multiple cognitive domains of attention, working memory, and positive or negative valence is the feature-based reward value learning task we used. This task can entail sub-conditions that quantify reversal learning flexibility (when the objects stay the same across blocks and only the reward contingencies change), as well as intra- and extras-dimensional set shifting abilities which are widely used markers of executive functioning (Crofts et al., 1999; Weed et al., 2008; Buckley et al., 2009; Wright et al., 2013; Shnitko et al., 2017; Azimi et al., 2020) with a high translational value (Keeler and Robbins, 2011). Here, we tested a feature-based version of reward learning rather than on object- or space-based learning because feature specific learning is considered the key learning strategy in naturalistic environments where even simple objects are composed of two or more dimensions (Farashahi et al., 2017; Womelsdorf et al., 2020). The results with this task may therefore prove to have high face validity about the real-world cognitive flexibility of subjects.

Similar to the visual search task, the feature-based reward learning task is easily extended to include other RDoC Matrix constructs such as loss aversion and the sensitivity of subjects to the positive and negative valence of outcomes (Evans et al., 2012; Banaie Boroujeni et al., 2020). For example, using visual tokens as secondary rewards we recently showed with the KS-1 that monkeys in some situations learned faster in the feature-based task when they could earn more tokens for correct choices but slowed down when they were losing tokens they already possessed (Banaie Boroujeni et al., 2020). The influence of prospective token-gains and token-losses measures the sensitivity of subjects to the valence of feedback which is one of five major domains of the RDoC Matrix (Cuthbert and Insel, 2013). In addition to varying the two tasks we described here, there are multiple further extensions conceivable. Previous work with rhesus monkeys in cage-based touchscreen settings showed that these task variations can reliably measure working memory, perceptual classification or transitive inferences, amongst others (Fagot and Paleressompoulle, 2009; Gazes et al., 2013; Hutsell and Banks, 2015; Calapai et al., 2017; Curry et al., 2017; Fizet et al., 2017; Berger et al., 2018; Sacchetti et al., 2021). Such cognitive testing is not restricted to rhesus monkeys as prior work showed cognitive engagement with touchscreens in multiple species including baboons (Fagot and Paleressompoulle, 2009; Fagot and De Lillo, 2011; Rodriguez et al., 2011; Claidiere et al., 2017), capuchin monkeys (Evans et al., 2008), marmosets (Kangas et al., 2016; Walker et al., 2020), and others (Hopkins et al., 1996; Beran et al., 2005; Egelkamp and Ross, 2019).

### Components integrated in KS-1

The successful use of KS-1 is not based on novel individual components but on the novel combination of components that allowed integrating it fully in the daily routines of the vivarium (**appendix 5.1**). There are four primary components that we consider particularly noteworthy. Firstly, its modular design enables the same Kiosk to be mounted to differently sized apartment cages by using custom tailored Kiosk frames (**Fig. 1A**). This feature enables using the Kiosk with different cage systems. Secondly, the KS-1 hosts the touchscreen, the computer hardware and user interface in a compact, closed cabinet inside the animal housing which reduces the outgoing cable to only the power line. The integrated cabinet enables using it in spaces that do not offer external spaces and it minimizes strain from un- and reconnecting cables (Calapai et al., 2017). Thirdly, KS-1 follows an open-source policy for all custom designed components with documentation and manuals for the major technical components (see **appendix 5.2**). This is a crucial feature designed to facilitate adoption of the behavioral enrichment and assessment tool in other contexts, closely following the tenets of OpenBehavior.com (White et al., 2019). Fourthly, the integration of behavioral control with a video engine designed for 3D rendered computer gaming (unity3D) enables conceiving of naturalistic task settings and virtual reality renderings that are not easily achieved by existing behavioral control software. However, we should note that KS-1 can be operated with other behavioral control software, for which many have been used in cage-based contexts, including control suites based on LabView (Grant et al., 2008; Shnitko et al., 2017), matlab (The Mathworks) using monkeylogic (Hwang et al., 2019; Sacchetti et al., 2021) or custom scripts (Griggs et al., 2021), C++ libraries as in MWorks (Calapai et al., 2017; Berger et al., 2018) (https://mworks.github.io), Presentation (Kret et al., 2016), Java (Fizet et al., 2017), Microsoft Visual Basic (Micheletta et al., 2015), Inquisit (McGuire et al., 2017), or E-Prime (Fagot and Paleressompoulle, 2009; Allritz et al., 2016). Through our adoption of *USE* in KS-1 we hope to not only provide a freely accessible software suite that is temporally precise and fully integrated with the KS-1 hardware systems (Watson et al., 2019b), but to inspire the development of naturalistic computer-game like tasks that can engage animals as well as humans and have been documented to be more motivating than simpler tasks (Bennett et al., 2016).

In summary, we outlined a cage-mounted kiosk station system that is integrated into the regular housing routines of an NHP vivarium, is highly engaging for animals, straightforward to operate by personnel, and rich in opportunities to discover cognitive capacities and strategies of NHP’s over prolonged periods of time.

## 5. Appendix

### 5.1. Overview of Kiosk hardware components

The Kiosk consists of multiple parts whose assembly is described in a manual available on the repository at https://github.com/att-circ-contrl/KioskStation. The following surveys the hardware components and computer related equipment of the KS-1:

- Kiosk Front-End (Stainless steel + Aluminum with frame for standard apartment cage (here: from Primate Products, Inc.) and sipper tube) and Kiosk back-end cabinet serving as electronics enclosure with three shelving levels and four 120mm 34 dBA DC brushless fans. An initial design of both parts is available in a cad file (see ≤https://github.com/att-circ-contrl/KioskStation>);
- Behavioral control PC machine (for running unity3D *USE* experimental suite and connecting to WIFI for data transfer) (the various models we used include: Intel NUC series NUC6i7KYK and NUC8i7HVK with 250GB SATA drives);
- Multi-camera control network ‘Neurocam’. The Neurocam includes a computer (NUC6i7), up to five cameras, and a LED strobe system, WIFI router, and a tablet for controlling camera streaming and monitoring the animals remotely. It is documented and available at https://github.com/att-circ-contrl/NeuroCam. For adjusting or rebuilding the firmware a user needs the NeurAVR library (https://github.com/att-circ-contrl/NeurAVR);
- I/O Synchbox for transferring TTL signals and synchronizing devices. The I/O Synchbox system is documented online with open-sourced firmware at ≤https://github.com/att-circ-contrl/SynchBox>. For adjusting and rebuilding the firmware the user needs the NeurAVR library (https://github.com/att-circ-contrl/NeurAVR);
- User interface with small monitor (LCD Eyoyo 12” with 1366×768p), keyboard (DAAZEE Ultra-Slim Small78 Keys Keyboard), and trackball mouse;
- LED Touchscreen (19.5” ELO Open Frame, 2094L);
- Industry grade fluid-reward pump system (LMI A741-910SI) with custom mount;
- Pellet dispenser (med-associates inc. 190mg Dispenser Pedestal) with custom mount;
- Foot-Operated Mobile Scissor-Style Lift Table Cart (enabling a single person to connect and disconnect Kiosk from apartment cage).

### 5.2. Online resources

Resources needed to build, assemble, and operate the KS-1 kiosk system for testing, training and enriching NHP’s are available online. The resources for the KS-1, the I/O Synchbox, the NeuroCam, and the USE behavioral control suite are available in github repositories (at https://github.com/att-circ-contrl/), via the Attention-Circuits-Control laboratory at http://accl.psy.vanderbilt.edu/kiosk/, or by contacting the corresponding author of this article. The resources include technical drawings of an initial Kiosk design (see **Fig. 1A**), technical drawings for mounting adaptors, a manual with multiple photos and technical details for assembling the Kiosk, and guides about how-to install and use the NeuroCam, the I/O Synchbox, and the remote triggering ‘reward’ fobs. The resources also include an installation guide, manual and example behavioral control configuration for the USE behavioral control suite (via https://github.com/att-circ-contrl/USE or <http://accl.psy.vanderbilt.edu/resources/analysis-tools/unifiedsuiteforexperiments/>).

## Acknowledgements

The authors thank Kevin Barker and Adrian Mizell for indispensable help with the kiosk design, and Shelby Volden and Seth König for help with initial animal training and Quaddle generation.

## Notes

### Competing Interest Statement

The authors have declared no competing interest.

## References

Allritz M, Call J, Borkenau P (2016) How chimpanzees (Pan troglodytes) perform in a modified emotional Stroop task. Anim Cogn 19:435–449.

Azimi M, Oemisch M, Womelsdorf T (2020) Dissociation of nicotinic alpha7 and alpha4/beta2 sub-receptor agonists for enhancing learning and attentional filtering in nonhuman primates. Psychopharmacology (Berl) 237:997–1010.

Bala PC, Eisenreich BR, Yoo SBM, Hayden BY, Park HS, Zimmermann J (2020) Automated markerless pose estimation in freely moving macaques with OpenMonkeyStudio. Nat Commun 11:4560.

Banaie Boroujeni K, Watson MR, Womelsdorf T (2020) Gains and Losses Differentially Regulate Attentional Efficacy and Learning at Low and High Attentional Load. bioRxiv https://doi.org/10.1101/2020.09.01.278168:1-38.

Bennett AJ, Perkins CM, Tenpas PD, Reinebach AL, Pierre PJ (2016) Moving evidence into practice: cost analysis and assessment of macaques’ sustained behavioral engagement with videogames and foraging devices. Am J Primatol 78:1250–1264.

Beran MJ, Beran MM, Harris EH, Washburn DA (2005) Ordinal judgments and summation of nonvisible sets of food items by two chimpanzees and a rhesus macaque. J Exp Psychol Anim Behav Process 31:351–362.

Berger M, Calapai A, Stephan V, Niessing M, Burchardt L, Gail A, Treue S (2017) Standardized automated training of rhesus monkeys for neuroscience research in their housing environment. J Neurophysiol:jn 00614 02017.

Berger M, Calapai A, Stephan V, Niessing M, Burchardt L, Gail A, Treue S (2018) Standardized automated training of rhesus monkeys for neuroscience research in their housing environment. J Neurophysiol 119:796–807.

Brainard DH (1997) The Psychophysics Toolbox. Spat Vis 10:433–436.

Buchanan RW, Keefe RS, Umbricht D, Green MF, Laughren T, Marder SR (2011) The FDA-NIMH-MATRICS guidelines for clinical trial design of cognitive-enhancing drugs: what do we know 5 years later? Schizophrenia bulletin 37:1209–1217.

Buchanan RW, Davis M, Goff D, Green MF, Keefe RS, Leon AC, Nuechterlein KH, Laughren T, Levin R, Stover E, Fenton W, Marder SR (2005) A summary of the FDA-NIMH-MATRICS workshop on clinical trial design for neurocognitive drugs for schizophrenia. Schizophrenia bulletin 31:5–19.

Buckley MJ, Mansouri FA, Hoda H, Mahboubi M, Browning PG, Kwok SC, Phillips A, Tanaka K (2009) Dissociable components of rule-guided behavior depend on distinct medial and prefrontal regions. Science 325:52–58.

Butler JL, Kennerley SW (2018) Mymou: A low-cost, wireless touchscreen system for automated training of nonhuman primates. Behav Res Methods.

Calapai A, Berger M, Niessing M, Heisig K, Brockhausen R, Treue S, Gail A (2017) A cage-based training, cognitive testing and enrichment system optimized for rhesus macaques in neuroscience research. Behav Res Methods 49:35–45.

Claidiere N, Gullstrand J, Latouche A, Fagot J (2017) Using Automated Learning Devices for Monkeys (ALDM) to study social networks. Behav Res Methods 49:24–34.

Crofts HS, Muggleton NG, Bowditch AP, Pearce PC, Nutt DJ, Scott EA (1999) Home cage presentation of complex discrimination tasks to marmosets and rhesus monkeys. Lab Anim 33:207–214.

Curry MD, Zimmermann A, Parsa M, Dehaqani M-RA, Clark KL, Noudoost B (2017) A Cage-Based Training System for Non-Human Primates. AIMS Neuroscience 4:102–119.

Cuthbert BN, Insel TR (2013) Toward the future of psychiatric diagnosis: the seven pillars of RDoC. BMC Med 11:126.

Datta SR, Anderson DJ, Branson K, Perona P, Leifer A (2019) Computational Neuroethology: A Call to Action. Neuron 104:11–24.

Doucet G, Gulli RA, Martinez-Trujillo JC (2016) Cross-species 3D virtual reality toolbox for visual and cognitive experiments. J Neurosci Methods 266:84–93.

Eastman KM, Huk AC (2012) PLDAPS: A Hardware Architecture and Software Toolbox for Neurophysiology Requiring Complex Visual Stimuli and Online Behavioral Control. Front Neuroinform 6:1.

Egelkamp CL, Ross SR (2019) A review of zoo_Jbased cognitive research using touchscreen interfaces. Zoo biology 38:220–235.

Evans TA, Beran MJ, Paglieri F, Addessi E (2012) Delaying gratification for food and tokens in capuchin monkeys (Cebus apella) and chimpanzees (Pan troglodytes): when quantity is salient, symbolic stimuli do not improve performance. Anim Cogn 15:539–548.

Evans TA, Beran MJ, Chan B, Klein ED, Menzel CR (2008) An efficient computerized testing method for the capuchin monkey (Cebus apella): adaptation of the LRC-CTS to a socially housed nonhuman primate species. Behav Res Methods 40:590–596.

Fagot J, Paleressompoulle D (2009) Automatic testing of cognitive performance in baboons maintained in social groups. Behav Res Methods 41:396–404.

Fagot J, Bonte E (2010) Automated testing of cognitive performance in monkeys: use of a battery of computerized test systems by a troop of semi-free-ranging baboons (Papio papio). Behav Res Methods 42:507–516.

Fagot J, De Lillo C (2011) A comparative study of working memory: immediate serial spatial recall in baboons (Papio papio) and humans. Neuropsychologia 49:3870–3880.

Farashahi S, Rowe K, Aslami Z, Lee D, Soltani A (2017) Feature-based learning improves adaptability without compromising precision. Nat Commun 8:1768.

Fizet J, Rimele A, Pebayle T, Cassel JC, Kelche C, Meunier H (2017) An autonomous, automated and mobile device to concurrently assess several cognitive functions in group-living non-human primates. Neurobiology of learning and memory 145:45–58.

Gazes RP, Brown EK, Basile BM, Hampton RR (2013) Automated cognitive testing of monkeys in social groups yields results comparable to individual laboratory-based testing. Anim Cogn 16:445–458.

Grant KA, Leng X, Green HL, Szeliga KT, Rogers LS, Gonzales SW (2008) Drinking typography established by scheduled induction predicts chronic heavy drinking in a monkey model of ethanol self-administration. Alcohol Clin Exp Res 32:1824–1838.

Griggs DJ, Bloch J, Chavan S, Coubrough KM, Conley W, Morrisroe K, Yazdan-Shahmorad A (2021) Autonomous cage-side system for remote training of non-human primates. Journal of Neuroscience Methods 348.

Harrison JE, Lophaven S, Olsen CK (2016) Which Cognitive Domains are Improved by Treatment with Vortioxetine? Int J Neuropsychopharmacol.

Hassani SA, Oemisch M, Balcarras M, Westendorff S, Ardid S, van der Meer MA, Tiesinga P, Womelsdorf T (2017) A computational psychiatry approach identifies how alpha-2A noradrenergic agonist Guanfacine affects feature-based reinforcement learning in the macaque. Sci Rep 7:40606.

Hopkins WD, Washburn DA, Hyatt CW (1996) Video-task acquisition in rhesus monkeys (Macaca mulatta) and chimpanzees (Pan troglodytes): a comparative analysis. Primates 37:197–206.

Hutsell BA, Banks ML (2015) Effects of environmental and pharmacological manipulations on a novel delayed nonmatching-to-sample ‘working memory’ procedure in unrestrained rhesus monkeys. J Neurosci Methods 251:62–71.

Hwang J, Mitz AR, Murray EA (2019) NIMH MonkeyLogic: Behavioral control and data acquisition in MATLAB. J Neurosci Methods 323:13–21.

Jacob G, Katti H, Cherian T, Das J, Zhivago KA, Arun SP (2020) A naturalistic environment to study natural social behaviors and cognitive tasks in freely moving monkeys. bioRxiv 311555:1–52.

Kangas BD, Bergman J, Coyle JT (2016) Touchscreen assays of learning, response inhibition, and motivation in the marmoset (Callithrix jacchus). Anim Cogn 19:673–677.

Karashchuk P (2019) Anipose. GitHub repository. https://github.com/lambdaloop/anipose. In.

Keeler JF, Robbins TW (2011) Translating cognition from animals to humans. Biochem Pharmacol 81:1356–1366.

Knight MJ, Baune BT (2018) Cognitive dysfunction in major depressive disorder. Curr Opin Psychiatry 31:26–31.

Krakauer JW, Ghazanfar AA, Gomez-Marin A, MacIver MA, Poeppel D (2017) Neuroscience Needs Behavior: Correcting a Reductionist Bias. Neuron 93:480–490.

Kret ME, Jaasma L, Bionda T, Wijnen JG (2016) Bonobos (Pan paniscus) show an attentional bias toward conspecifics’ emotions. Proc Natl Acad Sci U S A 113:3761–3766.

Lepora NF, Pezzulo G (2015) Embodied choice: how action influences perceptual decision making. PLoS Comput Biol 11:e1004110.

Mandell DJ, Sackett GP (2008) A computer touch screen system and training procedure for use with primate infants: Results from pigtail monkeys (Macaca nemestrina). Dev Psychobiol 50:160–170.

Mathis A, Mamidanna P, Cury KM, Abe T, Murthy VN, Mathis MW, Bethge M (2018) DeepLabCut: markerless pose estimation of user-defined body parts with deep learning. Nat Neurosci 21:1281–1289.

McGuire MC, Vonk J, Johnson-Ulrich Z (2017) Ambiguous Results When Using the Ambiguous-Cue Paradigm to Assess Learning and Cognitive Bias in Gorillas and a Black Bear. Behav Sci (Basel) 7.

Micheletta J, Whitehouse J, Parr LA, Waller BM (2015) Facial expression recognition in crested macaques (Macaca nigra). Anim Cogn 18:985–990.

Nagahara AH, Bernot T, Tuszynski MH (2010) Age-related cognitive deficits in rhesus monkeys mirror human deficits on an automated test battery. Neurobiol Aging 31:1020–1031.

National-Research-Council (1998) The Psychological Well-Being of Nonhuman Primates. Washington, DC: The National Academies Press.

Nuechterlein KH, Barch DM, Gold JM, Goldberg TE, Green MF, Heaton RK (2004) Identification of separable cognitive factors in schizophrenia. Schizophrenia research 72:29–39.

Peirce JW (2008) Generating Stimuli for Neuroscience Using PsychoPy. Front Neuroinform 2:10.

Perdue BM, Beran MJ, Washburn DA (2018) A computerized testing system for primates: Cognition, welfare, and the Rumbaughx. Behav Processes 156:37–50.

Rodriguez JS, Zurcher NR, Bartlett TQ, Nathanielsz PW, Nijland MJ (2011) CANTAB delayed matching to sample task performance in juvenile baboons. J Neurosci Methods 196:258–263.

Rumbaugh DM, Richardson WK, Washburn DA, Savage-Rumbaugh ES, Hopkins WD (1989) Rhesus monkeys (Macaca mulatta), video tasks, and implications for stimulus-response spatial contiguity. J Comp Psychol 103:32–38.

Sacchetti S, Ceccarelli F, Ferrucci L, Benozzo D, Brunamonti E, Nougaret S, Genovesio A (2021) Macaque monkeys learn and perform a non-match-to-goal task using an automated home cage training procedure. Scientific Reports 11:1–13.

Sheshadri S, Dann D, Hueser T, Scherberger H (2020) 3d reconstruction toolbox for behavior tracked with multiple cameras. Journal of Open Source Software 5:1849.

Shnitko TA, Allen DC, Gonzales SW, Walter NA, Grant KA (2017) Ranking Cognitive Flexibility in a Group Setting of Rhesus Monkeys with a Set-Shifting Procedure. Front Behav Neurosci 11:55.

Slifkin AB, Newell KM (1998) Is variability in human performance a reflection of system noise? Current Directions in Psychological Science 7:170–176.

Truppa V, Garofoli D, Castorina G, Piano Mortari E, Natale F, Visalberghi E (2010) Identity concept learning in matching-to-sample tasks by tufted capuchin monkeys (Cebus apella). Anim Cogn 13:835–848.

Tulip J, Zimmermann JB, Farningham D, Jackson A (2017) An automated system for positive reinforcement training of group-housed macaque monkeys at breeding and research facilities. J Neurosci Methods 285:6–18.

Unity-Technologies (2019) In.

Walker JD, Pirschel F, Gidmark N, MacLean JN, Hatsopoulos NG (2020) A platform for semiautomated voluntary training of common marmosets for behavioral neuroscience. J Neurophysiol 123:1420–1426.

Washburn DA, Rumbaugh DM (1992) Investigations of rhesus monkey video-task performance: evidence for enrichment. Contemp Top Lab Anim Sci 31:6–10.

Washburn DA, Hopkins WD, Rumbaugh DM (1991) Perceived control in rhesus monkeys (Macaca mulatta): enhanced video-task performance. J Exp Psychol Anim Behav Process 17:123–129.

Watson MR, Voloh B, Naghizadeh M, Womelsdorf T (2019a) Quaddles: A multidimensional 3-D object set with parametrically controlled and customizable features. Behav Res Methods 51:2522–2532.

Watson MR, Voloh B, Thomas C, Hasan A, Womelsdorf T (2019b) USE: An integrative suite for temporally-precise psychophysical experiments in virtual environments for human, nonhuman, and artificially intelligent agents. J Neurosci Methods 326:108374.

Weed MR, Bryant R, Perry S (2008) Cognitive development in macaques: attentional set-shifting in juvenile and adult rhesus monkeys. Neuroscience 157:22–28.

Weed MR, Taffe MA, Polis I, Roberts AC, Robbins TW, Koob GF, Bloom FE, Gold LH (1999) Performance norms for a rhesus monkey neuropsychological testing battery: acquisition and long-term performance. Brain Res Cogn Brain Res 8:185–201.

White SR, Amarante LM, Kravitz AV, Laubach M (2019) The Future Is Open: Open-Source Tools for Behavioral Neuroscience Research. eNeuro 6.

Wolfe JM (2021) Guided Search 6.0: An updated model of visual search. Psychon Bull Rev.

Wolfe JM, Horowitz TS (2017) Five factors that guide attention in visual search. Nature Human Behavior 1:1–8.

Womelsdorf T, Watson MR, Tiesinga P (2020) Learning at variable attentional load requires cooperation between working memory, meta-learning and attention-augmented reinforcement learning. bioRxiv https://doi.org/10.1101/2020.09.27.315432:1-50.

Wright MJ, Jr., Vandewater SA, Parsons LH, Taffe MA (2013) Delta(9)Tetrahydrocannabinol impairs reversal learning but not extra-dimensional shifts in rhesus macaques. Neuroscience 235:51–58.

Zhu Y, Womer FY, Leng H, Chang M, Yin Z, Wei Y, Zhou Q, Fu S, Deng X, Lv J, Song Y, Ma Y, Sun X, Bao J, Wei S, Jiang X, Tan S, Tang Y, Wang F (2019) The Relationship Between Cognitive Dysfunction and Symptom Dimensions Across Schizophrenia, Bipolar Disorder, and Major Depressive Disorder. Front Psychiatry 10:253.

